# Shifts in spring migration phenology are greater in northern wintering species

**DOI:** 10.1101/2025.03.24.644893

**Authors:** Tatsuya Miyamoto, Michio Kondoh

**Affiliations:** Graduate School of Life Sciences, Tohoku University; Graduate School of Life Sciences, Tohoku University, Advanced Institute for Marine Ecosystem Change, Tohoku University & JAMSTEC

## Abstract

Climate change is significantly affecting animal distributions and phenology, with migratory birds experiencing notable shifts in migration timing. There are complexities to changes in migration timing, considering variations due to monitoring locations or definitions of migration timing. Previous studies have focused on differences between traits in the magnitude of change and agreed that migration distance has an effect, although distribution has not been considered. This study investigates the factors influencing phenological changes in spring migration, focusing on First Arrival Date (FAD) and Last Departure Date (LDD) in eastern Japan. Using community science data from Sendai and Tokyo, we examine the effects of species characteristics, including trophic level, migration distance, and breeding and wintering areas, in addition to climatic variables. Consistent with other studies, FAD was advanced and LDD was delayed with climate. Notably, the magnitude of change in FAD differed depending on the distribution of wintering areas. Birds use many areas during their life cycle, and our results suggest that distribution, as a simple indicator, may be key to predicting responses to climate change.

## Introduction

Rapid climate change in recent years is causing major changes in the distribution and phenology of animals (Ambrosini et al., 2019; Chen et al., 2011). Changes in the timing of migration, possibly due to climate change, have been reported for a wide range of taxa, including mammals, fish, and insects (Hickling et al., 2006). For migratory birds, long-term observations have shown that the timing of migration is changing for various species in many regions (Cotton, 2003). Migratory birds in the northern hemisphere have two main migration periods per year: spring (pre-nuptial) migration to breeding sites and autumn (post-nuptial) migration to wintering sites. Most previous studies have reported an advance in spring arrival and a delay in autumn departure (Sparks et al., 2007).

Changes in migration timing can be detected with variations depending on the monitoring location on the migration route and the definition of migration timing, even in the same season and for the same species. Regarding differences in monitoring location, earlier departures do not necessarily result in earlier arrivals at the breeding sites. In fact, tracking studies of long-distance migration using geolocators show that birds modulate the timing of arrival at the breeding sites by adjusting the number of days spent at stopover sites (Conklin et al., 2021). This means that the phenological change is less detectable close to the breeding site. Modulation at the stopover site also affects the difference in the definition of migration timing. Representative methods for measuring migration timing are First Arrival Date (FAD) and Last Departure Date (LDD), indicating the first and the last observed day of the season, respectively. Fewer studies have been conducted on the LDD in spring than on FAD (Beaumont et al., 2006; Sparks et al., 2007).

Many researchers have examined species traits as factors explaining interspecific variations in the magnitude of changes in migration timing (Ambrosini et al., 2019). However, there is not always agreement among studies on the effects of traits; one study found that large, molting feathers before departure, and insects, seeds, or fruit-eating species were more likely to migrate later in the autumn, in which the timing of molt was the most important (Bitterlin & Josh, 2014). In contrast, another study reported that foraging guild, population trend, and relative abundance had no significant effect on changes in migration timing (Hurlbert & Liang, 2012). This discrepancy between studies may be due to the large intraspecific variation in the effects of climate, which varies with geographic location (Beissinger & Riddell, 2021).

Migratory birds move between several regions in an annual cycle, using each region for different purposes such as breeding, stopover, and wintering. Therefore, climatic changes in the different regions along the route can influence phenological changes. For example, birds wintering in cold regions with snow or frost are likely to be more dynamically affected by climate change (Bunn et al., 2007). Therefore, understanding changes in the timing of migration at a particular (observation) location would need to consider both climate variation at that site and the effects of other regions.

While it is challenging to account for all factors along the migration route, species distribution is a candidate for a simple indicator to consider regional differences. In this study, we use bird observation data in eastern Japan (Sendai and Tokyo) to understand which factors contribute to phenological changes in FAD and LDD in spring migration. The effects of species distribution are confirmed by including trophic level, migratory distance, and breeding and wintering region in the model, in addition to climatic effects. The data used in this study were obtained as community science, which are more suited to considering the phenology of multi-species over a migration period than using survey data of limited duration and species.

## Method

### Bird data

We used observation datasets from Sendai and Tokyo in eastern Japan. In Sendai, Tohoku University Bird Watching Club conducted weekly observations along the fixed route spanning 4 km from 2012 to 2022. For Tokyo, records from Tokyo, Kanagawa, and Saitama prefectures were obtained from eBird (Sullivan et al., 2009). The data period extends from 2009 to 2022.

The spring FAD and LDD for each year and species were obtained. The initial date was set as 1st January of each year, and the first record up to 70 was designated as the FAD, with the date of the last record up to 135 being set as the LDD. This straightforward definition enables the acquisition of FAD and LDD for all species while automatically excluding resident birds. To ensure consistent survey intervals, the data were summarized every five days.

### Climatic data

Two climatic data sets were utilized to consider environmental effects on phenological change. The North Pacific Index (NPI), measured based on atmospheric pressure, was obtained from the National Oceanic and Atmospheric Administration (Zhang et al., 1997). Monthly average and minimum temperatures in Sendai and Tokyo obtained from the Japan Meteorological Agency. Migration timings are thought to determine depending on the climate (especially minimum temperature) before the migration season (Redlisiak et al., 2018), so that we averaged each variables from February to April. Furthermore, the roll mean temperature and NPI from February to April over the past three years were utilized, as it is hypothesized that each individual bird selects the optimal migration time according to the past climate.

### Species trait

We investigated whether the magnitude of change in FAD and LDD differed depending on the difference in species traits considering four ordinal variables: trophic level, migration distance, northernmost wintering region, and southernmost breeding region. Trophic levels were categorized as carnivorous, omnivorous, and herbivorous based on the previous study (Pigot et al., 2020). Migration distance (3 categories), northernmost region of wintering (4 categories), and southernmost region of breeding (4 and 3 for FAD and LDD, respectively) were defined based on the distribution obtained from the Birds of the World (Billerman et al., 2022). For southernmost breeding region, a category “south” means south of Japan in FAD and south of Taiwan in LDD due to regional biases of observed species.

### Statistical analysis

We used a generalized linear mixed model (GLMM) in package lme4 (Bates et al., 2015) to test whether FAD and LDD could be explained by climate data, species traits, and the interaction between climate and traits. In the model, species were included as random intercepts to account for species-specific variability. Variable selection was based on the Akaike Information Criterion using the MuMIn package (Bartoń, 2024) and the number of used traits was limited to one or less.

## Result

The spring FAD was advanced 7.3 days per decade for 36 species (Figure 1). For example, FAD of Barn Swallow (*Hirundo rustica*) and Asian House-Martin (*Delichon dasypus*) were about 11 days earlier during the data period. In contrast, spring LDD was delayed by 10.6 days per decade for 63 species. Based on the variable selection process, the variation in the FAD was explained by the average temperature, NPI, with a significant interaction observed with wintering distribution (Table 1). The magnitude of the change was greater in the species that wintering in relatively northern region. In contrast, LDD showed no significant interactions with traits and was explained by mean and minimum temperatures (Table 1). In addition, LDD was potentially early on mid-distance migration species which does not show contribution to the magnitude of phenological change (Table 2).

**Table 1.**
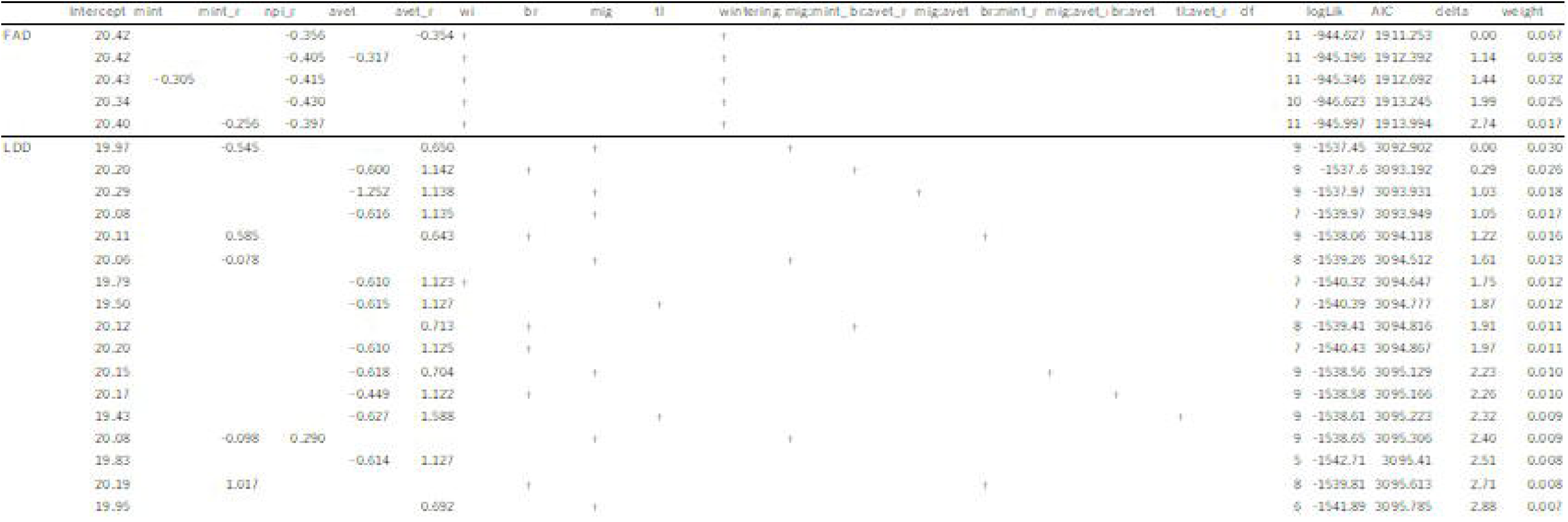
Result of variable selection. Arranged in order of explanation likelihood.

**Table 2.** Fixed effects of the best model. Mint: minimum temperature, avet: average temperature, r: roll mean, wi: northernmost wintering region, br: southernmost breeding region, mig: migration distance, tl: trophic level.

|  |  | Estimate | SE | t |
| --- | --- | --- | --- | --- |
| FAD | (Intercept) | 20.41587 | 1.366138 | 14.944 |
|  | avet_r | -0.35392 | 0.143497 | -2.466 |
|  | npi_r | -0.35564 | 0.451254 | -0.788 |
|  | wi(Honshu) | 2.188572 | 1.857535 | 1.178 |
|  | wi(SEA) | 4.252866 | 1.580502 | 2.691 |
|  | wi(Taiwan) | 2.723615 | 1.486607 | 1.832 |
|  | wi(Honshu):npi_r | -2.63196 | 0.6963 | -3.78 |
|  | wi(SEA):npi_r | -0.03127 | 0.502979 | -0.062 |
|  | wi(Taiwan):npi_r | -0.00371 | 0.483634 | -0.008 |
| LDD | (Intercept) | 21.2011 | 0.7186 | 29.502 |
|  | avet_r | 1.7795 | 0.258 | 6.898 |
|  | mint_r | -1.3523 | 0.2392 | -5.653 |
|  | mig(mid) | -2.426 | 0.933 | -2.6 |
|  | mig(short) | -1.133 | 1.1104 | -1.02 |

**Figure 1.**
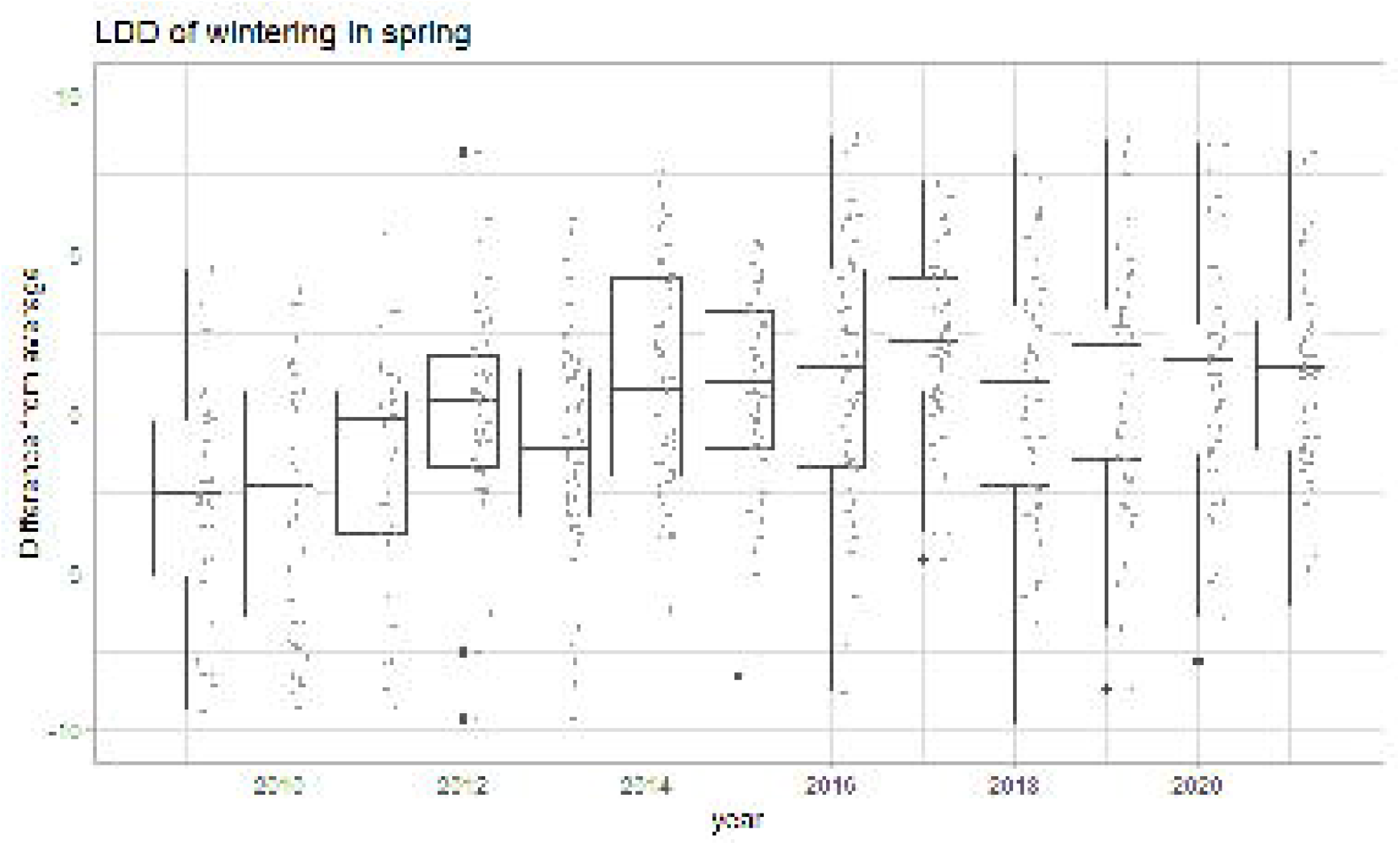
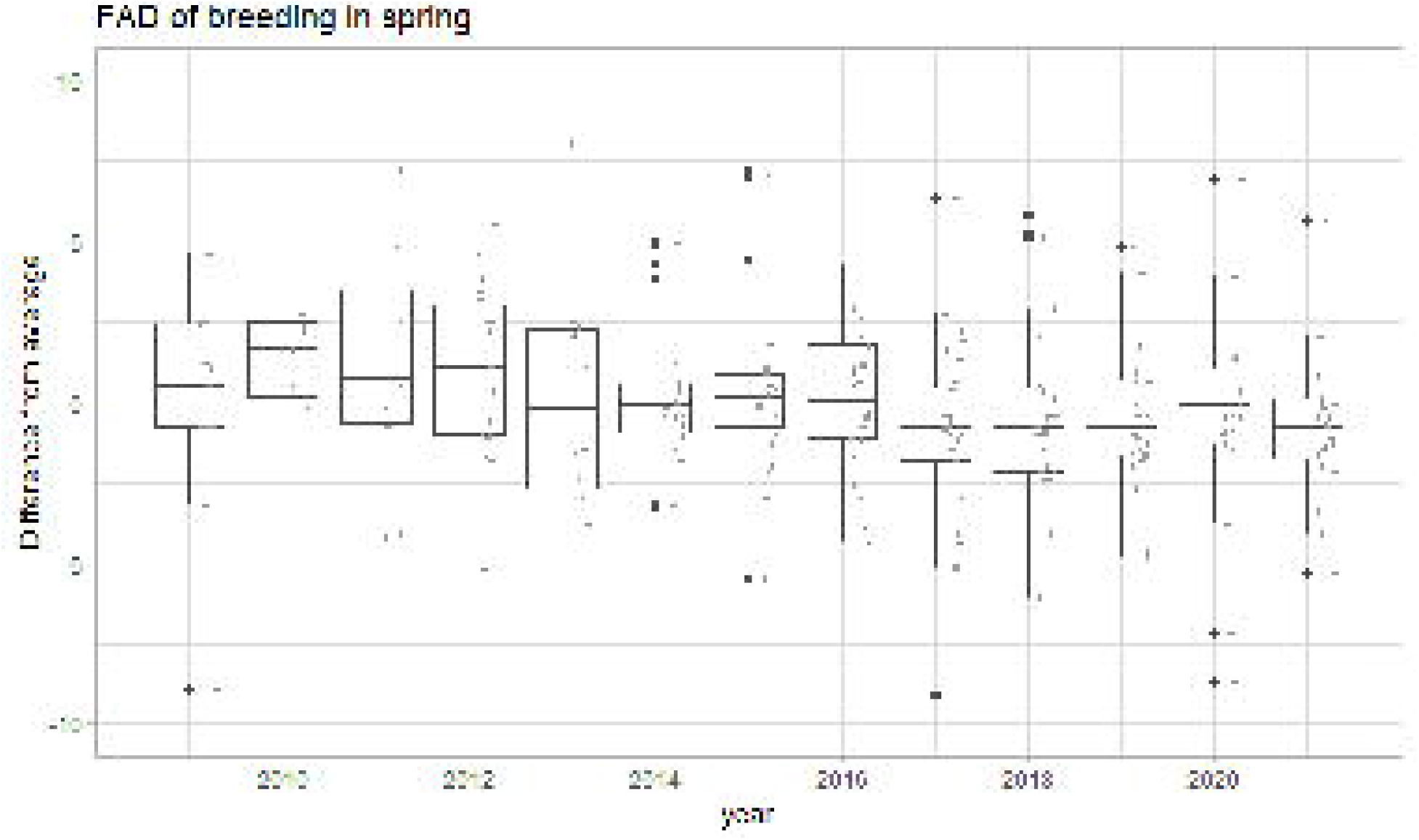
Overall trends of spring FAD and LDD. The average of each species was set to zero.

## Discussion

Bird records from two regions of Japan showed that the timing of spring migration is changing. The FAD was advanced 7.3 days per decade, which is notably higher than the previously reported range of 1.4 to 2.9 days per decade (Usui et al., 2017). However, it should be noted that the comparability of the speed of change in FAD is limited by variations in target species and long-term trends. Nonetheless, there is a possibility that the shift in FAD could have occurred over a brief period, exhibiting a marked acceleration in change. In terms of climate, the higher the temperature of the observation site, the more advanced the FAD and the more delayed the LDD. Furthermore, an elevated NPI has been demonstrated to induce early FAD. The NPI, an indicator of broad-area climate, exhibits fluctuations over several years; however, it demonstrated a consistent increase during the study period. This phenomenon may provide a rationale for the accelerated rate of change observed in FAD as compared to earlier studies, which employed more protracted data collection periods.

The observed differences in the magnitude of phenological change among species were best explained by the wintering locations. This finding underscores the necessity of incorporating distribution into the analysis of species-specific phenological variation. Two potential explanations for the impact of wintering grounds on observed phenological responses are proposed. The former is attributed to climatic variations between sites (departure hypothesis), while the latter is associated with the geographical distance between the observation site and the wintering location (stop-over hypothesis). **Departure hypothesis**: The phenomenon of climate change has been demonstrated to exert a greater influence on higher latitudes than on lower latitudes with respect to avian wintering patterns. For instance, waterfowl such as ducks necessitate unfrozen water surfaces for rest and foraging. In contrast, ground-foraging birds such as thrushes are limited in their foraging due to snow cover. These survival-critical thresholds of freezing and snowing are more likely to occur at high latitudes, and high-latitude ecosystems are affected by greater climate change (Bunn et al., 2007). A similar phenomenon can be expected in low-latitude regions, where small changes in weather conditions can significantly impact avian phenology. **Stop-over hypothesis**: In the case of a greater distance between the wintering site and the observation site, it is possible that some birds will remain at stop-over sites for a greater duration and in greater numbers. Consequently, the impact of the wintering site becomes less evident when the distance from the wintering site to the observation site is considerable. The absence of location interactions at LDD may be attributable to temporal adjustment by the climate at the stopover site, thereby nullifying the effect during the overwinter period. This phenomenon is further substantiated by prior research, which identified migration speed as the most significant factor (Hurlbert & Liang, 2012). It is plausible that species with slow migration patterns remain at the stopover site for a longer duration, potentially offsetting the effects of climate change at their wintering location.

Many studies have indicated that short-distance migratory species exhibit a greater degree of rapid change in relation to long-distance migrators concerning species-specific variations in the magnitude of observed phenological change (Buskirk et al., n.d.; Hubálek & Čapek, 2008). For instance, some farmland migrants have earlier arrival dates in western Poland (Tryjanowski et al., 2002). However, the significance of location over distance is a likely factor. In this study, migratory distance and the location of wintering and breeding sites were considered, but the model selection process identified the location of the wintering site as the most explanatory. In past studies, migratory distance has been a subject of discussion due to the finding that, if a high number of birds are breeding at an observation site, those with longer migratory distances are more likely to winter at lower latitudes, which can be explained by the same trend as location. Earlier studies on phenological variation have been conducted in Europe; however, the effects can be isolated when regional bias is reduced (Wormworth et al., 2006). A notable study demonstrated that the timing of changes varied according to the distribution of wintering populations (Maggini et al., 2020). This finding stands in contrast to the prevailing trend of migratory distance, thereby underscoring the notion that distributional variations play a major role in elucidating the phenological patterns exhibited by diverse species.

The study considered three categories of trophic level. Herbivorous species exhibited larger changes than carnivorous and omnivorous species; however, no stronger trend was identified than wintering location. While numerous traits have been examined in prior studies, there is a paucity of consensus on trends, with the exception of those related to migration and comparisons among species that winter in the same location. These findings offer a valuable contribution to our understanding of the impact of traits on the timing of migratory patterns (Ambrosini et al., 2019).

This study has an imprecation in conservation. Changes in migration timing have the potential to exert a deleterious effect on populations, given their capacity to engender temporal or spatial mismatches (Both et al., 2009). For instance, given the historical adaptability of migration timing, its shift can result in a reduction of food availability during the breeding season, thereby diminishing the reproductive success of the population (Visser & Gienapp, 2019). The importance of considering wintering locations, as outlined in this study, underscores the necessity for management and prioritization of diverse wintering site populations. The development of winter atlases or trans-seasonal observations has emerged as a valuable tool for understanding species-level and population-level shifts. These data, supported by local communities, facilitate the monitoring of species distribution and population dynamics over time. The enhancement of community observations, which provide multi-site and multi-temporal records, would facilitate the monitoring of phenology and population dynamics within a region.

## Supporting information

Supplemental Informations

## Acknowledgement

We thank the Tohoku University Birdwatching Club for providing us with bird observation data.

