## Supplemental Informations for "Shifts in spring migration phenology are greater in northern wintering species"

Supporting information 1. Bird traits.

| species | TrophicLevel | mig | wi | br(FAD) | br(LDD) |
| --- | --- | --- | --- | --- | --- |
| Cygnus cygnus | Herbivore | mid | hokkaido | okhotsk | okhotsk |
| Aix galericulata | Herbivore | short | honshu | china | south |
| Anas platyrhynchos | Herbivore | mid | okhotsk | japan | japan |
| Anas acuta | Herbivore | long | hokkaido | okhotsk | okhotsk |
| Anas crecca | Herbivore | long | hokkaido | okhotsk | okhotsk |
| Aythya ferina | Herbivore | long | okhotsk | okhotsk | okhotsk |
| Aythya fuligula | Omnivore | mid | okhotsk | okhotsk | okhotsk |
| Aythya marila | Omnivore | long | hokkaido | okhotsk | okhotsk |
| Bucephala clangula | Carnivore | mid | okhotsk | okhotsk | okhotsk |
| Mergellus albellus | Carnivore | long | hokkaido | okhotsk | okhotsk |
| Mergus merganser | Carnivore | mid | hokkaido | okhotsk | okhotsk |
| Mergus serrator | Carnivore | mid | hokkaido | okhotsk | okhotsk |
| Apus pacificus | Carnivore | long | SEA | china | south |
| Cuculus poliocephalus | Carnivore | long | taiwan | china | south |
| Cuculus optatus | Carnivore | long | SEA | japan | japan |
| Cuculus canorus | Carnivore | long | SEA | china | south |
| Treron sieboldii | Herbivore | short | honshu | no | south |
| Rallus indicus | Carnivore | mid | honshu | japan | japan |
| Gallinula chloropus | Omnivore | short | honshu | no | south |
| Fulica atra | Herbivore | short | honshu | no | south |
| Tachybaptus ruficollis | Carnivore | short | hokkaido | no | south |
| Podiceps grisegena | Carnivore | mid | okhotsk | okhotsk | okhotsk |
| Podiceps cristatus | Carnivore | mid | honshu | japan | japan |
| Podiceps auritus | Carnivore | mid | hokkaido | okhotsk | okhotsk |
| Vanellus vanellus | Carnivore | mid | honshu | japan | japan |
| Thinornis placidus | Carnivore | short | honshu | japan | japan |
| Thinornis dubius | Carnivore | long | taiwan | no | south |
| Actitis hypoleucos | Carnivore | mid | honshu | japan | japan |
| Tringa brevipes | Carnivore | long | SEA | okhotsk | okhotsk |
| Chroicocephalus ridibundus | Carnivore | mid | hokkaido | okhotsk | okhotsk |
| Larus crassirostris | Carnivore | short | okhotsk | china | south |
| Larus canus | Carnivore | long | okhotsk | okhotsk | okhotsk |
| Larus vegae | Carnivore | long | honshu | okhotsk | okhotsk |
| Phalacrocorax carbo | Carnivore | short | honshu | china | south |
| Nycticorax nycticorax | Carnivore | no | honshu | no | south |
| Butorides striata | Carnivore | short | taiwan | china | south |
| Ardea cinerea | Carnivore | short | hokkaido | no | south |
| Ardea alba | Carnivore | short | honshu | no | south |
| Pandion haliaetus | Carnivore | mid | hokkaido | no | south |
| Accipiter nisus | Carnivore | mid | okhotsk | japan | japan |
| Buteo japonicus | Carnivore | mid | okhotsk | japan | japan |
| Ninox japonica | Carnivore | long | taiwan | china | south |
| Alcedo atthis | Carnivore | short | honshu | no | south |
| Dendrocopos major | Omnivore | no | okhotsk | china | south |
| Falco tinnunculus | Carnivore | short | hokkaido | no | south |
| Pericrocotus divaricatus | Carnivore | long | SEA | japan | japan |
| Terpsiphone atrocaudata | Carnivore | long | taiwan | china | south |
| Garrulus glandarius | Omnivore | no | okhotsk | no | south |
| Cyanopica cyanus | Omnivore | no | hokkaido | japan | japan |
| Corvus frugilegus | Omnivore | short | honshu | japan | japan |
| Bombycilla garrulus | Herbivore | mid | hokkaido | okhotsk | okhotsk |
| Bombycilla japonica | Omnivore | long | hokkaido | okhotsk | okhotsk |
| Periparus ater | Omnivore | short | okhotsk | china | south |
| Poecile montanus | Omnivore | no | okhotsk | japan | japan |
| Hirundo rustica | Carnivore | long | taiwan | china | south |
| Delichon dasypus | Carnivore | long | taiwan | china | south |
| Cecropis daurica | Carnivore | short | taiwan | china | south |
| Urosphena squameiceps | Carnivore | long | taiwan | japan | japan |
| Phylloscopus coronatus | Carnivore | long | SEA | japan | japan |
| Phylloscopus borealoides | Carnivore | long | taiwan | japan | japan |
| Phylloscopus examinandus | Carnivore | long | SEA | okhotsk | okhotsk |
| Acrocephalus orientalis | Carnivore | long | SEA | japan | japan |
| Regulus regulus | Carnivore | short | hokkaido | japan | japan |
| Troglodytes troglodytes | Carnivore | no | hokkaido | china | south |
| Spodiopsar cineraceus | Carnivore | short | honshu | japan | japan |
| Agropsar philippensis | Omnivore | long | taiwan | japan | japan |
| Zoothera aurea | Carnivore | mid | honshu | japan | japan |
| Turdus cardis | Omnivore | long | taiwan | japan | japan |
| Turdus pallidus | Omnivore | mid | honshu | japan | japan |
| Turdus chrysolaus | Omnivore | short | honshu | japan | japan |
| Turdus eunomus | Omnivore | long | hokkaido | okhotsk | okhotsk |
| Muscicapa dauurica | Carnivore | long | taiwan | japan | japan |
| Cyanoptila cyanomelana | Carnivore | long | taiwan | japan | japan |
| Larvivora akahige | Carnivore | long | taiwan | japan | japan |
| Ficedula narcissina | Carnivore | long | SEA | japan | japan |
| Tarsiger cyanurus | Carnivore | short | honshu | japan | japan |
| Phoenicurus auroreus | Omnivore | mid | honshu | okhotsk | okhotsk |
| Monticola solitarius | Carnivore | short | hokkaido | china | south |
| Saxicola stejnegeri | Carnivore | long | taiwan | japan | japan |
| Cinclus pallasii | Carnivore | no | okhotsk | no | south |
| Passer cinnamomeus | Omnivore | mid | honshu | japan | japan |
| Prunella rubida | Carnivore | no | okhotsk | japan | japan |
| Motacilla cinerea | Carnivore | mid | honshu | china | south |
| Anthus hodgsoni | Carnivore | mid | honshu | japan | japan |
| Anthus japonicus | Carnivore | long | hokkaido | okhotsk | okhotsk |
| Fringilla montifringilla | Herbivore | long | hokkaido | okhotsk | okhotsk |
| Coccothraustes coccothraustes | Herbivore | mid | honshu | okhotsk | okhotsk |
| Eophona personata | Herbivore | short | honshu | china | south |
| Pyrrhula pyrrhula | Herbivore | short | okhotsk | japan | japan |
| Carpodacus sibiricus | Herbivore | mid | honshu | okhotsk | okhotsk |
| Spinus spinus | Herbivore | mid | hokkaido | okhotsk | okhotsk |
| Emberiza fucata | Herbivore | mid | honshu | japan | japan |
| Emberiza rustica | Herbivore | long | hokkaido | okhotsk | okhotsk |
| Emberiza elegans | Omnivore | mid | honshu | china | south |
| Emberiza personata | Carnivore | mid | honshu | china | south |
| Emberiza variabilis | Herbivore | short | honshu | japan | japan |

Supporting information 2. FAD.

| site | species | 2012 | 2013 | 2014 | 2015 | 2016 | 2017 | 2018 | 2019 | 2020 | 2021 | 2022 |
| --- | --- | --- | --- | --- | --- | --- | --- | --- | --- | --- | --- | --- |
| Sendai | Ardea cinerea | 19 | NA | NA | NA | 23 | NA | NA | 25 | 21 | 18 | NA |
| Sendai | Delichon dasypus | 20 | 21 | 19 | NA | 23 | NA | 20 | 17 | 17 | 18 | 16 |
| Sendai | Phylloscopus borealoides | 26 | 23 | 24 | NA | NA | NA | 24 | NA | 24 | 24 | 23 |
| Sendai | Acrocephalus orientalis | 33 | 27 | NA | NA | NA | NA | NA | 29 | 28 | 26 | 27 |
| Sendai | Phalacrocorax carbo | 31 | 25 | NA | NA | 23 | NA | NA | NA | 15 | NA | NA |
| Sendai | Alcedo atthis | 21 | NA | 24 | NA | NA | 26 | 28 | NA | NA | NA | NA |
| Sendai | Ficedula narcissina | 23 | 23 | 24 | 25 | 25 | 26 | 24 | 24 | 23 | 24 | 22 |
| Sendai | Agropsar philippensis | 22 | 21 | 24 | 22 | 23 | 23 | 21 | 24 | 20 | 24 | 22 |
| Sendai | Phylloscopus coronatus | 26 | 22 | 24 | 22 | 23 | 21 | 21 | 23 | 23 | 22 | 21 |
| Sendai | Hirundo rustica | 19 | 20 | 18 | 18 | 22 | 20 | 18 | 17 | 15 | 18 | 18 |
| Sendai | Cuculus poliocephalus | 31 | 28 | 28 | 33 | NA | 31 | 31 | 29 | 28 | 29 | 26 |
| Sendai | Urosphena squameiceps | 31 | 22 | 24 | 22 | 23 | 21 | 23 | 22 | 23 | 23 | 22 |
| Sendai | Apus pacificus | NA | 24 | 26 | NA | NA | NA | NA | 24 | 23 | 23 | 24 |
| Sendai | Monticola solitarius | NA | 21 | NA | 19 | 25 | 20 | 23 | NA | 19 | NA | 30 |
| Sendai | Phylloscopus examinandus | NA | 29 | 31 | 32 | 32 | NA | 30 | 31 | 29 | 29 | 29 |
| Sendai | Cyanoptila cyanomelana | NA | 22 | 24 | 28 | 23 | 32 | 24 | 22 | 23 | 22 | 22 |
| Sendai | Motacilla cinerea | NA | 19 | 21 | NA | 26 | 19 | 24 | 18 | NA | NA | NA |
| Sendai | Turdus cardis | NA | 29 | 25 | NA | NA | NA | NA | NA | 24 | 22 | 20 |
| Sendai | Muscicapa dauurica | NA | 23 | 22 | NA | 23 | NA | 21 | 27 | 23 | 22 | 24 |
| Sendai | Anthus japonicus | NA | 15 | NA | NA | NA | NA | NA | NA | 17 | 21 | 18 |
| Sendai | Saxicola stejnegeri | NA | 20 | NA | NA | NA | NA | NA | 23 | 23 | 22 | 20 |
| Sendai | Bombycilla japonica | NA | 24 | 21 | NA | 22 | 21 | 20 | 23 | 20 | 18 | 18 |
| Sendai | Ninox japonica | NA | NA | 31 | NA | NA | NA | NA | 31 | 26 | 26 | 26 |
| Sendai | Terpsiphone atrocaudata | NA | NA | 31 | 32 | NA | 31 | 28 | 31 | 28 | 26 | 28 |
| Sendai | Emberiza fucata | NA | NA | 24 | 28 | 32 | NA | NA | NA | NA | NA | 21 |
| Sendai | Actitis hypoleucos | NA | NA | NA | 25 | 25 | NA | NA | NA | NA | 22 | 18 |
| Sendai | Pandion haliaetus | NA | NA | NA | 23 | 18 | NA | NA | NA | 19 | NA | 15 |
| Sendai | Spodiopsar cineraceus | NA | NA | NA | 21 | NA | NA | 27 | 18 | NA | 16 | 18 |
| Sendai | Thinornis dubius | NA | NA | NA | NA | NA | 21 | 21 | 20 | 18 | 18 | 17 |
| Sendai | Pericrocotus divaricatus | NA | NA | NA | NA | NA | NA | 24 | 25 | 23 | 23 | 23 |
| Tokyo | Apus pacificus | 21 | 19 | 24 | 30 | 22 | 25 | 28 | 18 | 19 | NA | NA |
| Tokyo | Delichon dasypus | 17 | 16 | NA | 22 | NA | 17 | NA | 16 | 15 | NA | NA |
| Tokyo | Acrocephalus orientalis | 24 | 24 | 23 | 24 | 19 | 23 | 23 | 23 | 23 | 23 | NA |
| Tokyo | Cyanoptila cyanomelana | 22 | NA | 22 | 21 | 20 | 23 | 21 | 19 | 23 | 22 | NA |
| Tokyo | Tringa brevipes | 27 | 27 | 23 | 24 | 24 | 24 | 24 | 24 | 23 | 23 | NA |
| Tokyo | Ficedula narcissina | 21 | 25 | 22 | 22 | 22 | 22 | 23 | 21 | 23 | 22 | NA |
| Tokyo | Turdus cardis | 21 | NA | 21 | 19 | 23 | 19 | 19 | 20 | 24 | 23 | NA |
| Tokyo | Cecropis daurica | 28 | NA | 32 | NA | 32 | 34 | NA | 23 | 19 | 26 | NA |
| Tokyo | Thinornis dubius | 16 | NA | NA | 22 | NA | NA | NA | NA | NA | NA | NA |
| Tokyo | Larvivora akahige | 27 | 21 | 24 | NA | 19 | 23 | 21 | 23 | NA | 29 | NA |
| Tokyo | Agropsar philippensis | 21 | 24 | NA | 22 | 23 | 21 | NA | 20 | 22 | 23 | NA |
| Tokyo | Phylloscopus coronatus | 28 | 21 | 22 | 23 | 22 | 21 | 23 | 21 | 22 | 22 | NA |
| Tokyo | Cuculus optatus | 24 | NA | 26 | NA | 25 | 25 | 24 | 29 | 24 | 23 | NA |
| Tokyo | Hirundo rustica | 20 | 15 | 16 | 18 | 19 | 16 | NA | 15 | NA | NA | NA |
| Tokyo | Cuculus poliocephalus | 29 | NA | 24 | 26 | 29 | 30 | 24 | 29 | 29 | 26 | NA |
| Tokyo | Urosphena squameiceps | 24 | NA | 22 | 20 | 23 | NA | 21 | NA | 21 | 22 | NA |
| Tokyo | Cuculus canorus | NA | 31 | NA | NA | 32 | 28 | 34 | NA | 28 | 29 | NA |
| Tokyo | Butorides striata | NA | 23 | NA | 33 | 23 | 25 | 27 | NA | 27 | 24 | NA |
| Tokyo | Phylloscopus borealoides | NA | NA | 23 | NA | 23 | 26 | 25 | 25 | 24 | 23 | NA |
| Tokyo | Pericrocotus divaricatus | NA | NA | 23 | NA | 26 | 24 | 23 | 23 | 23 | 20 | NA |
| Tokyo | Muscicapa dauurica | NA | NA | NA | 24 | NA | NA | 21 | 24 | 24 | 23 | NA |

Supporting information 3. LDD.

| site | species | 2013 | 2014 | 2015 | 2016 | 2017 | 2018 | 2019 | 2020 | 2021 | 2022 |
| --- | --- | --- | --- | --- | --- | --- | --- | --- | --- | --- | --- |
| Sendai | Emberiza personata | 27 | NA | 26 | NA | 24 | 27 | 27 | NA | NA | 27 |
| Sendai | Turdus chrysolaus | 15 | 24 | 25 | 25 | NA | 25 | 24 | 27 | NA | 27 |
| Sendai | Fringilla montifringilla | 23 | 17 | 8 | 19 | 16 | 16 | 21 | 25 | 8 | 19 |
| Sendai | Apus pacificus | 25 | NA | NA | NA | NA | NA | 24 | NA | 25 | 25 |
| Sendai | Eophona personata | 27 | 16 | NA | 8 | NA | NA | 27 | NA | 26 | NA |
| Sendai | Thinornis placidus | 24 | NA | NA | 12 | 27 | NA | NA | 24 | NA | NA |
| Sendai | Pyrrhula pyrrhula | 15 | 14 | 15 | NA | NA | NA | NA | 19 | NA | NA |
| Sendai | Phylloscopus borealoides | 27 | 24 | NA | NA | NA | 24 | NA | 27 | 27 | NA |
| Sendai | Cygnus cygnus | 7 | NA | 9 | NA | NA | NA | 10 | 11 | 9 | 12 |
| Sendai | Anas acuta | 14 | 14 | 25 | 11 | 26 | 17 | 14 | 20 | 9 | 15 |
| Sendai | Tachybaptus ruficollis | 11 | 11 | 9 | 15 | 17 | 18 | 20 | 21 | 18 | 23 |
| Sendai | Emberiza rustica | 18 | 19 | 14 | 19 | 7 | 14 | 18 | 10 | 19 | 23 |
| Sendai | Mergus merganser | 23 | 21 | NA | 22 | 26 | 21 | NA | 26 | 23 | NA |
| Sendai | Regulus regulus | 21 | 14 | 15 | 19 | 17 | NA | 20 | 21 | 9 | 14 |
| Sendai | Motacilla cinerea | 23 | 21 | 22 | NA | NA | NA | NA | NA | 26 | NA |
| Sendai | Aythya fuligula | 20 | 14 | 25 | 22 | 12 | 21 | 25 | 24 | 24 | 24 |
| Sendai | Emberiza variabilis | 27 | NA | NA | NA | NA | NA | NA | 25 | 26 | 27 |
| Sendai | Anas crecca | 25 | 26 | 26 | 25 | 27 | 25 | 27 | NA | NA | 26 |
| Sendai | Muscicapa dauurica | 25 | NA | NA | 23 | NA | 21 | 27 | NA | NA | NA |
| Sendai | Coccothraustes coccothraustes | 27 | 22 | 21 | 23 | 20 | 23 | 24 | NA | NA | NA |
| Sendai | Phoenicurus auroreus | 18 | 14 | 21 | 19 | 19 | 17 | 20 | 17 | 18 | 22 |
| Sendai | Turdus pallidus | 22 | 18 | 25 | 19 | 24 | 17 | 24 | 26 | 24 | 24 |
| Sendai | Phylloscopus coronatus | 27 | NA | NA | 26 | NA | NA | NA | 27 | 27 | NA |
| Sendai | Anthus japonicus | 15 | 22 | 8 | NA | NA | NA | NA | 24 | 24 | 22 |
| Sendai | Turdus eunomus | 25 | 25 | 23 | 25 | 26 | 24 | 24 | 25 | 25 | 25 |
| Sendai | Zoothera aurea | 11 | 22 | NA | 12 | NA | NA | 7 | NA | 25 | 12 |
| Sendai | Buteo japonicus | 24 | 11 | 18 | 18 | 21 | NA | 20 | NA | NA | NA |
| Sendai | Saxicola stejnegeri | 24 | NA | NA | NA | NA | 7 | 23 | 23 | 22 | 24 |
| Sendai | Accipiter nisus | 11 | 11 | NA | 8 | 17 | 20 | NA | 21 | 25 | 20 |
| Sendai | Periparus ater | 27 | 10 | 21 | 19 | NA | NA | 15 | 25 | 22 | 20 |
| Sendai | Bombycilla japonica | 24 | 24 | 23 | 22 | 24 | 23 | 23 | 24 | NA | 26 |
| Sendai | Carpodacus sibiricus | 19 | 18 | 15 | 19 | 17 | 18 | 17 | 24 | 18 | 22 |
| Sendai | Anas platyrhynchos | 24 | 18 | 25 | 19 | 23 | 16 | NA | 21 | 26 | 24 |
| Sendai | Spinus spinus | 27 | NA | 16 | 22 | 10 | 23 | NA | NA | NA | 23 |
| Sendai | Spodiopsar cineraceus | 22 | NA | 25 | NA | 24 | NA | 24 | NA | 27 | 27 |
| Sendai | Tarsiger cyanurus | 10 | NA | NA | 15 | 19 | 17 | 10 | NA | 22 | 20 |
| Sendai | Turdus cardis | NA | 25 | NA | NA | NA | NA | NA | 24 | 27 | 27 |
| Sendai | Agropsar philippensis | NA | 25 | 23 | NA | 26 | 27 | NA | NA | NA | NA |
| Sendai | Aythya ferina | NA | 14 | 14 | NA | 14 | NA | 15 | NA | 9 | NA |
| Sendai | Actitis hypoleucos | NA | NA | 26 | NA | 24 | 20 | 24 | NA | NA | 22 |
| Sendai | Ardea alba | NA | NA | 16 | NA | NA | 17 | NA | NA | 26 | NA |
| Sendai | Emberiza elegans | NA | NA | 7 | 13 | 12 | NA | 17 | NA | 11 | 13 |
| Sendai | Anthus hodgsoni | NA | NA | NA | 13 | NA | 16 | NA | 25 | 26 | 25 |
| Tokyo | Podiceps grisegena | 14 | 14 | NA | 22 | 19 | 22 | 14 | 14 | 19 | NA |
| Tokyo | Turdus chrysolaus | 25 | 23 | 22 | 26 | 24 | NA | 26 | 26 | 25 | NA |
| Tokyo | Fringilla montifringilla | 9 | 24 | 15 | 16 | NA | 23 | 25 | 11 | 21 | NA |
| Tokyo | Apus pacificus | 19 | 24 | NA | NA | 25 | NA | NA | NA | NA | NA |
| Tokyo | Actitis hypoleucos | 20 | 24 | NA | NA | NA | NA | NA | NA | NA | NA |
| Tokyo | Pyrrhula pyrrhula | 19 | 26 | 22 | 23 | 15 | 13 | 20 | 19 | 20 | NA |
| Tokyo | Mergus serrator | 16 | 17 | 24 | 13 | 23 | 23 | 22 | NA | 19 | NA |
| Tokyo | Larus schistisagus | 18 | NA | NA | NA | NA | NA | NA | NA | NA | NA |
| Tokyo | Fulica atra | 20 | 25 | 26 | NA | NA | NA | NA | NA | NA | NA |
| Tokyo | Aix galericulata | 17 | 17 | 15 | 22 | 20 | 23 | 17 | 16 | 17 | NA |
| Tokyo | Anas acuta | 20 | 24 | 18 | 23 | 27 | NA | 22 | 17 | 25 | NA |
| Tokyo | Emberiza rustica | 14 | 17 | 21 | 23 | 17 | 20 | 21 | 16 | 19 | NA |
| Tokyo | Larus canus | 16 | 24 | 24 | 21 | 27 | 21 | 20 | NA | 22 | NA |
| Tokyo | Podiceps cristatus | 16 | NA | NA | 24 | NA | NA | NA | NA | NA | NA |
| Tokyo | Regulus regulus | 21 | 26 | 22 | NA | 18 | 23 | 18 | NA | NA | NA |
| Tokyo | Aythya fuligula | 26 | NA | NA | NA | NA | NA | NA | NA | 22 | NA |
| Tokyo | Rallus indicus | 21 | 15 | 25 | 26 | 19 | 19 | NA | 21 | NA | NA |
| Tokyo | Emberiza variabilis | 24 | 25 | 23 | 21 | 20 | 19 | 23 | 21 | 23 | NA |
| Tokyo | Anas crecca | 22 | 25 | 26 | NA | NA | 25 | 27 | NA | NA | NA |
| Tokyo | Larvivora akahige | 21 | 24 | NA | 19 | 23 | 23 | 25 | 10 | NA | NA |
| Tokyo | Agropsar philippensis | 25 | NA | 22 | 23 | 24 | NA | NA | 23 | 26 | NA |
| Tokyo | Coccothraustes coccothraustes | 21 | 25 | 23 | 26 | 26 | 25 | 25 | NA | 26 | NA |
| Tokyo | Turdus pallidus | 24 | 24 | 23 | 23 | 23 | 23 | 25 | 23 | 23 | NA |
| Tokyo | Phoenicurus auroreus | 19 | 17 | 22 | 18 | 19 | 16 | 20 | 20 | 22 | NA |
| Tokyo | Anthus japonicus | 20 | 17 | 23 | 21 | 22 | 23 | 22 | 22 | 22 | NA |
| Tokyo | Falco tinnunculus | 15 | 26 | 23 | 26 | NA | NA | NA | NA | NA | NA |
| Tokyo | Turdus eunomus | 25 | 24 | 24 | 26 | 26 | 25 | 27 | 27 | 27 | NA |
| Tokyo | Bombycilla japonica | 21 | 18 | 18 | NA | NA | 23 | NA | 22 | 13 | NA |
| Tokyo | Anthus hodgsoni | 20 | 17 | NA | 25 | 26 | 14 | 23 | 23 | 19 | NA |
| Tokyo | Carpodacus sibiricus | 15 | 14 | 14 | 13 | 19 | 20 | 21 | 23 | 20 | NA |
| Tokyo | Aythya ferina | 18 | NA | NA | NA | NA | NA | NA | NA | 23 | NA |
| Tokyo | Spinus spinus | 19 | 16 | 22 | 16 | NA | 25 | 7 | 26 | 23 | NA |
| Tokyo | Pandion haliaetus | 12 | 19 | 12 | NA | NA | NA | NA | NA | 27 | NA |
| Tokyo | Corvus frugilegus | 18 | NA | NA | 26 | NA | 13 | NA | 14 | NA | NA |
| Tokyo | Chroicocephalus ridibundus | 26 | NA | 26 | NA | NA | NA | NA | NA | 26 | NA |
| Tokyo | Tarsiger cyanurus | 21 | 24 | 22 | 15 | 21 | 23 | 22 | 18 | 18 | NA |
| Tokyo | Phylloscopus borealoides | NA | 24 | NA | 26 | 26 | 25 | 25 | 26 | 27 | NA |
| Tokyo | Zoothera aurea | NA | 26 | 19 | NA | 24 | 21 | 13 | 26 | NA | NA |
| Tokyo | Buteo japonicus | NA | 14 | 21 | 15 | 25 | NA | NA | 19 | NA | NA |
| Tokyo | Accipiter nisus | NA | 13 | NA | NA | 19 | 19 | 21 | 19 | 23 | NA |
| Tokyo | Anas platyrhynchos | NA | 24 | 23 | NA | NA | NA | NA | NA | NA | NA |
| Tokyo | Podiceps auritus | NA | 15 | 12 | 22 | 16 | 14 | 24 | 13 | 11 | NA |
| Tokyo | Bombycilla garrulus | NA | NA | 18 | NA | NA | 13 | NA | 18 | 18 | NA |
| Tokyo | Vanellus vanellus | NA | NA | 11 | 15 | NA | 11 | 7 | 10 | 9 | NA |
| Tokyo | Bucephala clangula | NA | NA | 10 | 23 | 20 | 9 | 18 | 9 | 11 | NA |
| Tokyo | Emberiza elegans | NA | NA | 12 | 13 | 13 | NA | 7 | NA | NA | NA |
| Tokyo | Passer cinnamomeus | NA | NA | NA | 21 | 18 | NA | 17 | 15 | 14 | NA |
| Tokyo | Emberiza fucata | NA | NA | NA | 14 | 23 | NA | NA | 23 | 22 | NA |
| Tokyo | Mergellus albellus | NA | NA | NA | 14 | NA | 13 | 10 | 13 | 9 | NA |
